# An optimised method for bacterial nucleic acid extraction from positive blood culture broths for whole genome sequencing, resistance phenotype prediction and downstream molecular applications

**DOI:** 10.1101/2022.06.30.497578

**Authors:** Michelle J. Bauer, Anna Maria Peri, Lukas Lüftinger, Stephan Beisken, Haakon Bergh, Brian M. Forde, Cameron Buckley, Thom Cuddihy, Patrice Tan, David L. Paterson, David M. Whiley, Patrick N. A. Harris

## Abstract

**Background:** A prerequisite to rapid molecular detection of pathogens causing bloodstream infections is an efficient, cost effective and robust DNA extraction solution. We describe methods for microbial DNA extraction direct from positive blood culture broths, suitable for metagenomic sequencing and the application of machine-learning based tools to predict antimicrobial susceptibility.

**Methods:** Prospectively collected culture-positive blood culture broths with Gram-negative bacteria, were directly extracted using various commercially available kits. We compared methods for efficient inhibitor removal, avoidance of DNA shearing or degradation, to achieve DNA of high quality and purity. Bacterial species identified via whole-genome metagenomic sequencing (Illumina, MiniSeq) from blood culture extracts were compared to conventional methods from cultured isolates (Vitek MS). A machine-learning algorithm (AREScloud) was used to predict susceptibility against commercially available antibiotics, compared to susceptibility testing (Vitek 2) and other commercially available rapid diagnostic instruments (Accelerate Pheno and BCID).

**Results:** A two-kit method using a modified MolYsis Basic kit (for host DNA depletion) and extraction using Qiagen DNeasy UltraClean microbial kits resulted in optimal extractions appropriate for multiple molecular applications, including PCR, short-read and long-read sequencing. DNA extracts from 40 blood culture broths were included. Taxonomic profiling by direct metagenomic sequencing matched species identification by conventional methods in 38/40 (95%) of samples, with two showing agreement to genus level. In two polymicrobial samples, a second organism was missed by sequencing. Whole genome sequencing antimicrobial susceptibility testing (WGS-AST) models were able to accurately infer profiles for 6 common pathogens against 17 antibiotics. Overall categorical agreement (CA) was 95%, with 11% very major errors (VME) and 3.9% major errors (ME). CA for WGS-AST was >95% for 5/6 of the most common pathogens (*E. coli, K. pneumoniae, P. mirabilis, P. aeruginosa* and *C. jejuni*) while it was lower for *K. oxytoca* (66.7%), likely due to the presence of inducible cephalosporinases. Performance of WGS-AST was sub-optimal for uncommon pathogens (e.g. *Elizabethkingia*) and some combination antibiotic compounds (e.g. ticarcillin-clavulanate). Time to pathogen identification and resistance gene detection was fastest with BCID (1 h from blood culture positivity). Accelerate Pheno provided a rapid MIC result in approximately 8 h. While Illumina based direct metagenomic sequencing did not result in faster turn-around times compared conventional methods, use of real-time nanopore sequencing may allow faster data acquisition.

**Conclusions:** The application of direct metagenomic sequencing from positive blood culture broths is a feasible approach and solves some of the challenges of sequencing from low-bacterial load samples. Machine-learning based algorithms are also accurate for common pathogen / drug combinations, although additional work is required to optimise algorithms for uncommon species and more complex resistance genotypes, as well as streamlining methods to provide more rapid sequencing results.

## INTRODUCTION

Bloodstream infections are a major cause of morbidity and mortality, and rapid pathogen identification and antimicrobial susceptibility phenotyping is critical to patient outcome ^1,2^. Current pathogen identification and phenotypic antimicrobial susceptibility testing can take up to 3 days, or longer. Consequently, rapid molecular detection and gene profiling methodologies are desirable ^2-4^. There could be great benefit in a holistic feasible approach to attain a single host depleted DNA extraction direct from blood culture (BC) broth, suitable for multiple downstream molecular applications. Further, the implementation of commercial kits with minimal out-of-kit modifications, and potential automation, would ensure robust, reproducible results.

We aimed to develop a reliable DNA extraction method for direct whole genome sequencing from positive blood culture broths to support rapid bacterial characterisation and antibiotic susceptibility prediction in patients with bloodstream infections. A primary aim was to obtain human depleted and enriched microbial DNA extraction from blood culture broths suitable for multiple downstream molecular applications, including whole genome sequencing (WGS). The objective was to achieve high quality bacterial DNA extractions, depleted of human DNA and inhibitors (salts, proteins, enzymes, preservatives, neutralising compounds), of appropriate input length and at usable concentrations for WGS. Additionally, effects of bench top duration or freeze-thaw conditions were assessed. Downstream applications included traditional and real-time PCR, as well as short and long-read WGS using either enzyme based or ligation library preparation. As the ultimate goal of molecular testing from positive blood culture broth is clinical implementation ^5^, results obtained were assessed for their ability to predict clinically relevant microbial phenotypes (e.g. species identification, resistance gene detection and antibiotic susceptibility testing [AST]). To evaluate the ability to rapidly predict AST *in silico* direct from genomic data, we compared a machine-learning based whole genome sequencing AST (WGS-AST) tool to conventional culture-based methods. In addition, we compared these methods to commercially available rapid diagnostic systems based on morphokinetic cellular analysis and multiplex PCR.

## METHODS

### Blood cultures

Forty blood culture broths (FA plus, FN plus and paediatric PF plus bottles; bioMérieux) that flagged positive for mono or polymicrobial growth with Gram-negative bacteria (identified by Gram stain and microscopy) were included. These were collected from patients presenting to emergency departments or admitted to an intensive care unit (ICU) served by Central laboratory, Pathology Queensland, Brisbane, Australia. Two clinical samples containing CTX-M ESBL-producing *E coli* (DETECT-110 and DETECT-111) and one non-ESBL-producing *E. coli* (sample 112) were used for assay development and validation purposes. Two samples (DETECT-113, DETECT-114) had no positive growth and were used as negative controls. The subsequent 37 samples (DETECT-115 to DETECT-151) were tested prospectively across all testing platforms. Positive blood culture broths were de-identified and analysed at the University of Queensland, Centre for Clinical Research. Blood culture bottles were removed from BACT/Alert Virtuo System (bioMerieux) once flagged positive and were extracted by the research laboratory within 1.5 hours.

### Sample processing and storage

Blood cultures broths were extracted for genomic DNA upon receipt and the remaining sample frozen at -20 and -80⍰C, and if required, thawed to room temperature from frozen.

### Genomic DNA Extraction

Host genomic DNA (gDNA) was depleted using the MolYsis Complete, MolYsis Basic Kit 0.2mL and 1mL protocols (Molzyme, Germany), according to manufacturer’s instructions, with the following exceptions: The blood culture broth starting volume for the Basic kit was increased for higher yields from 0.2 to 0.5mL and manufacturer’s instructions followed. After the host depletion stage, samples were centrifuged at 10,000g for 30 seconds and the supernatant removed. The microbial pellet underwent gDNA extraction by one of the two methods (Method 1: “Mini-Pure” Extraction and Method 2: “UltraClean” Extraction; full details in Supplementary Material)

Genomic DNA quality and purity checks were undertaken by QUBIT fluorometer (Life Technologies), NanoDrop 2000 Spectrophotometer (Thermo Scientific) and Agilent TapeStation 4150 using Genomic DNA ScreenTape and Reagents or D1000 High Sensitivity for sequencing library preparations. In addition, we assessed the utility of gDNA size selection using the Circulomics Short Read Eliminator (Circulomics), inhibitor removal by QIAGEN DNeasy PowerClean Pro Clean up Kit and the NEBNext® Microbiome DNA Enrichment Kit Ethanol precipitation protocol modified with the removal of TE buffer and 12.5μL pre-warmed water for elution. Twenty Mini-pure (“Method 1”) extractions and twenty UltraClean extractions (“Method 2”) were prepared for short read sequencing (Figure 1).

**Figure 1:**
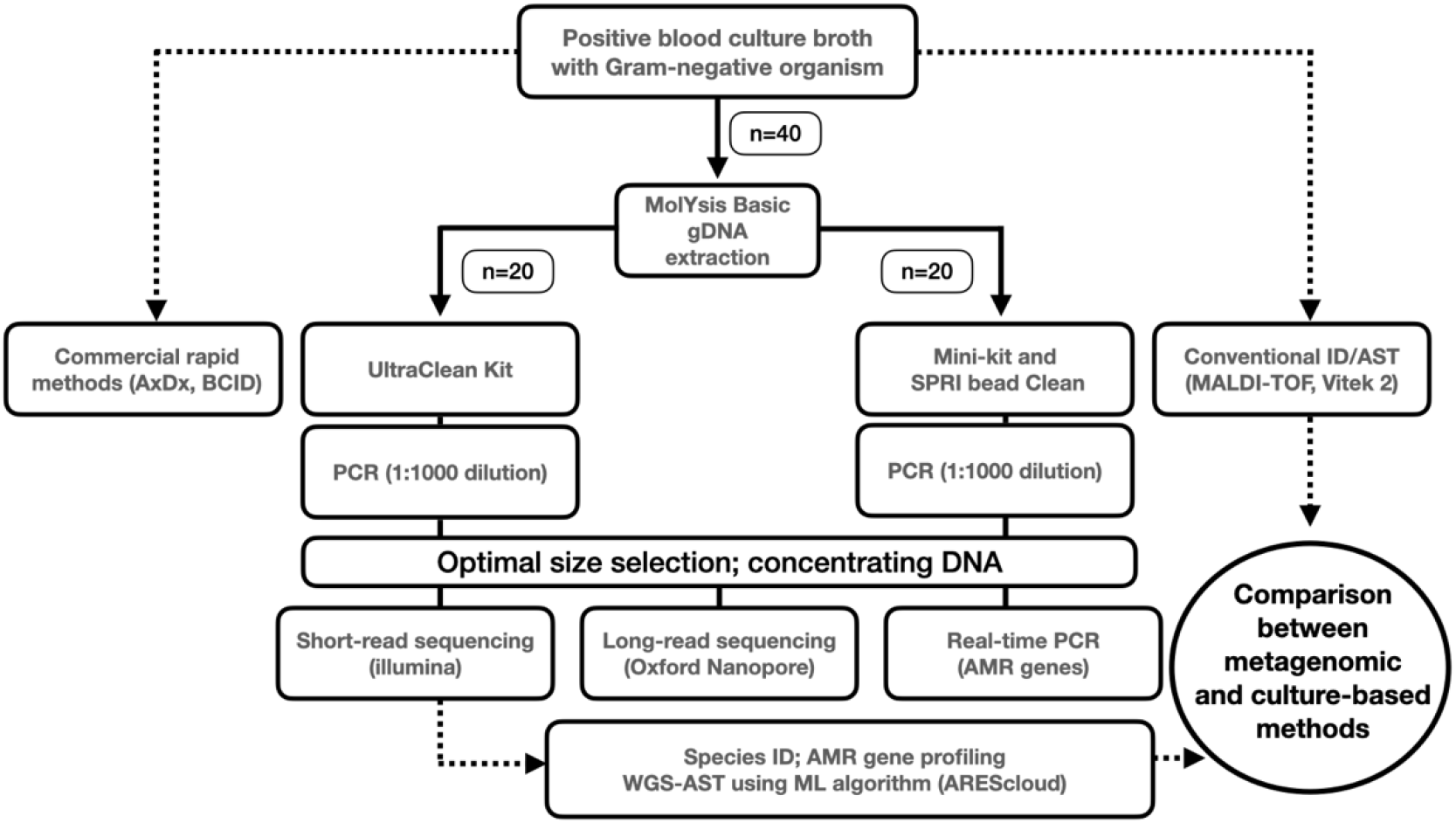
Study workflow. Direct metagenomic sequencing from positive blood culture broths compared with culture-based methods and commercial rapid diagnostics. Two DNA extraction methods were compared each using half (n=20) of the samples. AxDx = Accelerate Pheno; BCID = blood culture identification PCR panel (bioMérieux); gDNA = genomic DNA; SPRI = solid-phase reversible immobilization; PCR = polymerase chain reaction; ID = identification; AST = antimicrobial susceptibility testing; WGS = whole genome sequencing; MALDI-TOF = matrix-assisted laser desorption/ionization-time of flight; AMR = antimicrobial resistance; ML = machine learning

### Inhibitor removal and human genomic DNA depletion

Traditional and real-time PCR assays were undertaken to determine any effects of SPS being co-purified or acting as an inhibitor to PCR. Inhibition was assessed according to the positive control and experiment outcomes outlined by Regan *et al* ^6^. Dilutions were prepared from 1 in 10 to 1 in 10,000. There was no inhibition of PCR amplification as there were resolved bands of expected amplicon size for each dilution. This was observed with real-time PCR with cycle-to-threshold (Ct) values of 13 at 1 in 10 with 3-fold increases to a Ct of 27 for 1 in 10,000 dilutions (Supplementary Figure S1). TaqMan ERV-3 real-time PCR was undertaken to determine if the process of host human genomic DNA was depleted in the subsequent microbial DNA extraction. The assay confirmed complete human genomic DNA depletion (Supplementary Figure S2A) and 16S gene detected for the microbial DNA extraction (Supplementary Figure S2B).

### Sequencing

Short read sequencing from blood culture broth extractions utilised the Nextera DNA Flex Library Prep Kit (Illumina), with a modification of the starting input of 5 μL gDNA with samples >20 ng/μL. Pooled libraries were loaded in the Mid or High Output Reagent Cartridge (300 cycles) and sequenced as paired ends on the Illumina MiniSeq platform. Long read sequencing was undertaken with R9.4.1 flow cells as a singleplex using either the library preparation Rapid Sequencing or Ligation Kits, and multiplexing using the Rapid Barcoding kit or Ligation kit with Native Barcodes (Oxford Nanopore Technologies). All sequencing utilised the flow cell priming kit (EXP-FLP002) and voltage drift was accounted for where the flow cell went through a wash protocol.

### Metagenome assemblies, taxonomic profiling and WGS-AST using machine-learning

Assembly and binning for whole genome sequencing AST (WGS-AST) from metagenomes was performed using the pipeline nf-core/mag v2.1.1 ^7^. In short, raw reads were trimmed and mapped against the GRCh38 and PhiX genome to remove reads from contaminant species. Retained reads were assembled with both SPAdes ^8^ and MEGAHIT ^9^. Binning of assembled metagenomes into metagenomic bins was performed with MetaBAT2 ^9^. Taxonomy was assigned to metagenomic bins using GTDB-Tk ^10^. Completeness and duplication of bins was assessed with BUSCO ^11^ and QUAST ^12^. For each sample and assembly algorithm, taxonomy at the species-level could be assigned only to a single bin, with all remaining bins highly incomplete and likely not representing distinct pathogen species in the input sample. Downstream analysis was performed on whole metagenome assemblies. For each sample, the metagenome assembly (produced by either SPAdes or MEGAHIT) with the highest BUSCO completeness and lowest BUSCO duplication at domain level was selected. Selected metagenome assemblies were uploaded to the AREScloud web application, release 2022-01, (Ares Genetics GmbH, Vienna, AT) for genomic prediction of antimicrobial susceptibility. The platform used stacked classification machine learning (ML) WGS-AST models trained on ARESdb ^13^, combined with rule-based resistance prediction via ResFinder 4 ^14^ to provide species-specific susceptibility/resistance (S/R) predictions. AST predictions for a total of 17 antibiotic compounds were generated for samples belonging to six of the most common hospital-acquired pathogens. Very major error (VME) and major error (ME) rates were defined following CLSI M52 guidelines ^15^, and categorical agreement between results of WGS-AST and conventional AST were calculated for antimicrobial-organism combinations. *In silico* detection of resistance genes was determined by screening the genome assemblies for each isolate against the NCBI resistance gene database ^16,17^ using abricate v. 0.9.8 (https://github.com/tseemann/abricate) with default parameters.

### Real-time PCR for resistance genes

The utility of PCR for the targeted detection of key antimicrobial resistance (AMR) genes direct from blood culture extracts was also assessed using real-time TaqMan PCR assays. In brief, reaction mixes were prepared with 10 μL QuantiTect Probe PCR master mix (QIAGEN), 1.0 μM primer and 0.25 μM probe, 2.0 μL purified gDNA diluted 1 in 1000 in molecular grade water, with a total reaction volume of 20 μL. Assays included detection of ERV-3 for host gDNA depletion ^18^, 16S in microbial DNA extractions and the following antimicrobial resistance genes: 16S methylases [*armA, rmtF, rmtB, rmtC*] ^19^; extended-spectrum-β-lactamases (ESBLs) [*bla*_SHV 5/12_, *bla*_VEB_, *bla*_CTX-M_ group 1 & 9] ^20,21^; carbapenemases [*bla* _KPC_, *bla*_NDM_, *bla* _IMP-4_, *bla*_VIM_, *bla*_OXA-48-like_ ] ^22^; *ampC* [*bla*_CMY-2-like_ ] (this study, Supplementary Tables S1, S2); and colistin resistance [*mcr-1*] ^23,24^. Reactions were run on Rotor-Gene Q real-time PCR thermocycler (QIAGEN) 95⍰C for 15 minutes; 45 cycles at 95⍰C for 15 seconds, 60⍰C for 30 seconds. Result analysis was conducted with Rotor-Gene 6000 Series software. Sodium Polyanethole Sulfonate (SPS) removal assays were conducted by PCR gene amplification and gel electrophoresis to assess PCR inhibition of increasing 1 in 10 dilutions. GoTaq reaction mix was prepared with 6.5 μL GoTaq (Promega), 1.0 μM primers, 3.0 μL molecular grade water and 1 μL template. Cycling conditions (BioRad Thermocycler C1000 Touch) 95⍰C for 5 minutes; 35 cycles at 95⍰C for 40 seconds, 55⍰C for 1 minute and 72⍰C for 1 minute, 72⍰C for 5 minutes. A 3% agarose gel with 4 μL ethidium bromide was loaded with 5 μL PCR reaction and 2 μL GeneRuler 100bp ladder Plus (Thermo Fisher) and run for 40 minutes at 100 volts. Gel images were captured by ViberLourmat UV Gel Dock.

### Commercial Rapid Diagnostic Instruments

For comparison, positive blood culture broths were also tested using two commercially available platforms that provide rapid species identification and limited AMR gene profiling (Blood Culture Identification Panel [BCID] on the BioFire FilmArray Instrument; bioMérieux) as well as predictive MIC using morphokinetic cellular analysis (Accelerate Pheno; Accelerate Diagnostics). The BCID and Accelerate Pheno tests were performed as per manufacturer’s instructions, with the exemption of blood culture transfer to the BCID testing pouch by 27G x ½” needle (Henke Sass Wolf) and syringe.

### Conventional AST

All molecular, rapid diagnostic and genomic-based ID/AST testing was compared to conventional culture-based methods validated for clinical use at Pathology Queensland for diagnostic testing. Species identification was performed using MALDI-TOF (Vitek MS, bioMérieux) on pure cultured isolates, with AST performed by Vitek 2 automated broth microdilution (N-246 AST cards; bioMérieux), using EUCAST clinical breakpoints applicable at the time ^25^. For certain species (e.g. *Campylobacter jejuni*) AST was undertaken using disk diffusion according to EUCAST methods ^26^. Conventional testing was considered the standard against which molecular and genomic tests were compared.

### Ethics

Ethics was approved by the Royal Brisbane & Women’s Hospital and ratified by UQ Human Ethics Human Research Ethics Committee LNR/2018/QRBW/44671.

## RESULTS

### Blood Culture Microorganisms

Of the 37 positive blood culture broths, 35 had monomicrobial growth according to conventional ID methods, including *Escherichia coli* (n=14), *Klebsiella pneumoniae* (n=2) *and Klebsiella oxytoca* (n=1), *Enterobacter hormaechei* (n=1), *Morganella morganii* (n=1), *Proteus mirabilis* (n=3), *Pseudomonas aeruginosa* (n=5) and *Pseudomonas mosselii* (n=1), *Bacteroides thetaiotaomicron* (n=1), *Campylobacter jejuni* (n=2), *Elizabethkingia anophelis* (n=1), *Yokenella regensburgei* (n=1), *Pasteurella multocida* (n=1), *and Vogesella perlucida* (n=1). Two samples gave polymicrobial results with conventional culture testing (*E. coli* and *Enterococcus faecium*; *E. coli* and *K. pneumoniae* respectively) (Supplementary Data).

### DNA purity

DNA quality and purity were assessed from host depletion to subsequent microbial DNA extraction (Supplementary Table S3). The MolYsis Complete kit extraction process includes host depletion and microbial DNA extraction. Extraction concentrations were less than 1ng/μL with poor A_260_/A_230_ ratios, indicating potential extraction salt carryover. The MolYsis kit was consequently used for host depletion preceding microbial DNA extraction with the Mini or UltraClean kits. The Mini kit method had lower yields of DNA than the UltraClean method which was also of poor purity A_260_/A_230_ 0.6 and adequate A_260_/A_280_ 0.3 and low quantification DNA ratios. Purity was improved with SPRI bead cleanup which removed contaminants and or inhibitors and provided opportunity to increase concentration through low volume elution (Supplementary Table S3). An alternative method of ethanol precipitation was trialled to concentrate DNA; however, DNA was continually lost, and the method abandoned. Another process to remove inhibitors was through the QIAGEN DNeasy PowerClean column, purified DNA was eluted however there was a 50-80% loss in DNA concentration (results not shown). The UltraClean kit has an inhibitor removal reagent which is effective in purifying the DNA and with the amount of starting material, eluting high concentrations for downstream applications.

### DNA length, size selection and concentration

Depending on the downstream application, DNA length ranged from greater than 30Kbp with the Mini kit extraction to an average 16Kbp with the UltraClean kit (Supplementary Table S3). The application of the Circulomics Short Fragment Eliminator removed smaller fragments while maintaining input DNA concentration of longer fragments. Size selection with SPRI beads using ratio 1 in 1.5 removed smaller fragments of around 4Kbp without shearing longer fragments from either extraction method. To create smaller gDNA fragments, extractions were diluted 150, 100 and 50 μL and sheared for a fragment size of 10 Kbp, with resulting sizes from 11,048 to 11867 bp.

### Blood Culture Extractions under variable storage temperatures and duration

The viability of storage conditions for BC was assessed for concentration, purity, and length under two different storage conditions, room temperature for up to 5 days and thawed from frozen. DNA was directly extracted from BC by MolYsis and Mini Kit without the SPRI bead clean-up at day 1 and every 24 hours until day 5. The same BC sample was frozen at -20^°^C at day 1 and was thawed to room temperature on day 5 and extracted as previously described. Overall, DNA extraction concentration, purity and length of 2-5 days and frozen were comparable to the fresh BC baseline extraction (day 1).

### Downstream molecular applications

Forty direct BC extractions, twenty from each extraction method, that underwent short read sequencing, were analysed for taxonomy and predictive AST profiling, as well as real-time PCR for AMR gene detection. For extensive investigation of applications, one blood culture sample (DETECT-110; containing *E. coli* carrying *bla*_*CTX-M-15*_) was extracted by both methods along with the cultured isolate. Another blood culture sample (DETECT-112, containing non-ESBL *E. coli*) was used to assess time and temperature experiments and lastly, one additional blood culture was tested contained the reference *E. coli* strain EC958 with known plasmid number (Table 1). Short read sequencing was possible from both extraction methods and the AMR profiling confirmed 6 genes with the same identity; this was validated by real-time PCR for the *bla*_*CTX-M-15*_ as well as the expected absent genes. Identical AMR gene profiles were verified with long read sequencing by both extraction methods using the transposase library preparation, also observed with the UltraClean and ligation library preparation. The Mini-pure extraction with ligation library preparation and long read sequenced resulted in loss of genes. *E. coli* EC958 was sequenced using the UltraClean DNA extraction ^27^. With no size selection and both the transposase and ligation library prep, all plasmids were accounted for (Table 1).

**Table 1.** Downstream molecular applications: qPCR, short and long read sequencing from direct BC extractions and cultured isolates using two methods, including QC strain (EC958) and studies of effects of bench time and temperature

| Extraction | Molecular Application | Assay Type | AMR Gene detected | Taxonomy | Number Plasmids |
| --- | --- | --- | --- | --- | --- |
| **UltraClean:** EC958 isolate (QC strain) | Short Read | Nextera Flex | Yes | *E. coli* | 2 |
|  | Long Read | Transposase | Yes | *E. coli* | 2 |
|  |  | Ligation | Yes | *E. coli* | 2 |
| Extraction | Molecular Application | Assay Type | AMR Gene detected | Taxonomy | Number Genes<sup>d</sup> |
| **UltraClean:** BC containing CTX-M ESBL *E. coli*<sup>a</sup> | qPCR<sup>c</sup> | TaqMan Probe | Yes (Ct 22.0) | - | - |
|  | Short Read | Nextera Flex | Yes | *E. coli* | 6 |
|  | Long Read | Transposase | Yes | *E. coli* | 6 |
|  |  | Ligation | Yes | *E. coli* | 6 |
| **UltraClean:** BC containing CTX-M ESBL *E. coli*<sup>a</sup> | qPCR<sup>c</sup> | TaqMan Probe | Yes (Ct 23.0) | - | - |
|  | Short Read | Nextera Flex | Yes | *E. coli* | 6 |
|  | Long Read | Transposase | Yes | *E. coli* | 6 |
|  |  | Ligation | Yes | *E. coli* | 6 |
| **Mini-pure (+SPRI bead purify):** BC containing CTX-M ESBL *E. coli*<sup>a</sup> | qPCR<sup>c</sup> | TaqMan Probe | Yes (Ct 24.2) | - | - |
|  | Short Read | Nextera Flex | Yes | *E. coli* | 6 |
|  | Long Read | Transposase | Yes | *E. coli* | 6 |
|  |  | Ligation | Yes | *E. coli* | 5 |
| **Duration study (1-5 days):** using non-ESBL *E. coli*<sup>b</sup> | qPCR<sup>c</sup> | TaqMan Probe | no CTX-M to detect |  | - |
|  | Short Read | Nextera Flex | Yes | *E. coli* | 6 |
|  | Long Read | Transposase | Yes | *E. coli* | 6 |
| **Freeze-thaw:** using non-ESBL *E. coli*<sup>b</sup> | qPCR<sup>c</sup> | TaqMan Probe | no CTX-M to detect |  | - |
|  | Short Read | Nextera Flex | Yes | *E. coli* | 6 |
|  | Long Read | Transposase | Yes | *E. coli* | 6 |

<sup>a</sup> Sample DETECT-110: BC sample containing ESBL *E. coli* (*bla*<sub>CTX-M-15</sub>); <sup>b</sup> Sample DETECT-112: BC sample containing non-ESBL *E. coli*; <sup>c</sup> PCR to detect *bla*<sub>CTX-M-15</sub>; <sup>d</sup> DETECT-110 contained following AMR genes: *aadA5*, *bla*<sub>CTX-M-15</sub>, *bla*<sub>EC-5</sub>, *dfrA17*, *mph(A)*, *sul1*; DETECT-112 contained following AMR genes: *aph(3'')-Ib*, *aph(6)-Id*, *bla*<sub>EC-5</sub>, *bla*<sub>TEM-1</sub>, *dfrA5*, *sul2* [note *tet(34)* gene was found with only 73% coverage, so is not included in AMR gene number]

### Taxonomy identification of metagenomic samples and in silico predictive AST

Taxonomy identification of metagenomic samples down to species level yielded good agreement with conventional testing; for two samples identification to the genus level only was achieved, including *Vogesella urethralis* (which was identified by VITEK MS as *Vogesella perlucida*) and *Escherichia flexneri* (which was identified as by VITEK MS as *E. coli*). In two polymicrobial samples no presence of a second species was found during processing of metagenomic reads, with *E. coli* only being identified in each sample by metagenomic sequencing (Supplementary Data). Interestingly in one of the polymicrobial samples, the Accelerate Pheno and BCID2 systems detected the *E. faecium* but not the *E. coli*.

The performance of the whole genome sequencing AST (WGS-AST) models were assessed for a subset of 6 common pathogens and 17 antibiotic compounds. Overall categorical agreement (CA) was 95%, with 11% very major errors (VME; false prediction of susceptibility) and 3.9% major errors (ME; false prediction of resistance) (Table 2; Supplementary Table S4). CA was >95% for 5/6 of the common bloodstream pathogens (*E. coli, K. pneumoniae, P. mirabilis, P. aeruginosa* and *C. jejuni*) while it was lower for *K. oxytoca* (66.7%), reflecting errors in predicting ceftriaxone susceptibility, likely due to the challenge of chromosomally-encoded and inducible cephalosporinases such as *bla* _OXY-2_ ^*28*^.

**Table 2:** Performance of WGS-AST for 6 most common Gram-negative pathogens

| Taxon | Categorical Agreement (%) | VME (%) | ME (%) |
| --- | --- | --- | --- |
| All | 95 | 11 | 4 |
| *Escherichia coli* | 96 | 0 | 5 |
| *Klebsiella pneumoniae* | 96 |  | 4 |
| *Pseudomonas aeruginosa* | 97 | 0 | 3 |
| *Proteus mirabilis* | 100 | 0 | 0 |
| *Klebsiella oxytoca* | 67 | 67 | 0 |
| *Campylobacter jejuni* | 100 |  | 0 |

VME=very major error; ME = major error; blank cells reflect insufficient data

**Table 3:**
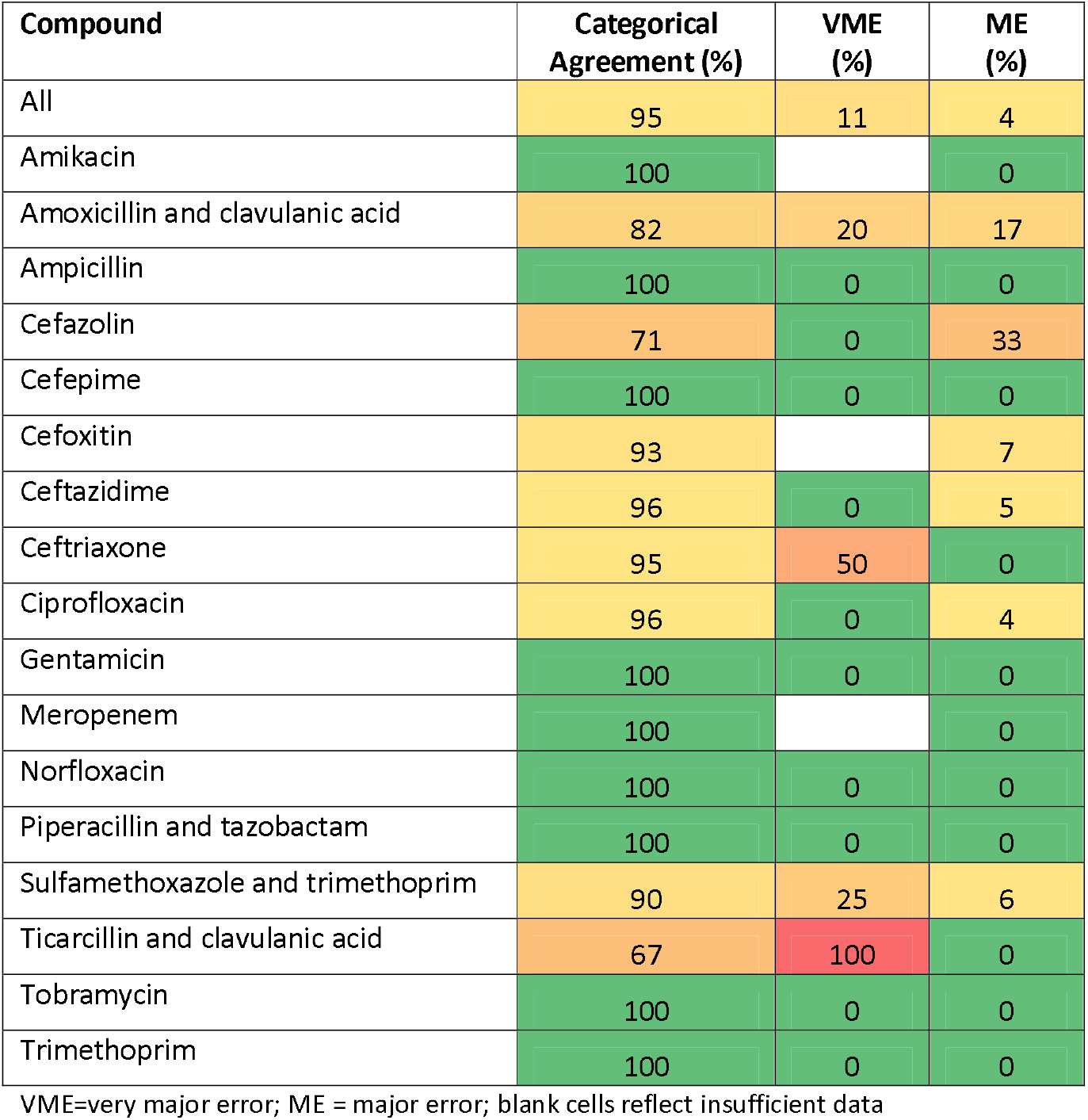
Performance of WGS-AST for Gram-negative active antibiotics across species tested

For exploratory research purposes, in cases where neither WGS-AST models nor species-relevant ResFinder 4 panels existed for uncommon pathogens, non-panel ResFinder 4 calls (based solely on presence of known AMR markers related to the compound in question, disregarding taxonomy) were used (Supplementary Table S5). The resulting set of WGS-AST calls encompassed all species found across metagenomic samples. Calls produced this way exhibited higher rates of very major error (VME) of 50.1%. This was particularly the case for taxa far removed from core ResFinder target taxa such as *Elizabethkingia* and *Yokenella*, and for combination agents such as ticarcillin/clavulanate. Specifically, out of 39 false susceptible exploratory predictions, 22 were for combination agents, and 13 were calls for unusual species for which no WGS-AST models nor species-relevant ResFinder 4 rules exist.

### Commercial Rapid Diagnostics

Thirty-seven samples were assessed with new rapid diagnostic commercially available platforms. Out of 35 monomicrobial samples, the Biofire Filmarray BCID2 panel was able to identify 4 pathogens at genus level and 22 at species level, although in two cases it gave an incorrect dual identification with *Proteus* being identified as a second genus in two samples harbouring *E. coli* and *K. pneumoniae* respectively. Similar performance was shown by the Accelerate Pheno which identified 7 pathogens at genus level and 18 at species level. In 8 cases the two instruments gave no ID and these all included off-panel pathogens; in one case only, a discordant result was observed (*Yokenella regensburgei* misidentified as *Enterobacter* spp.). Out of 2 polymicrobial samples both instruments correctly identified one of the 2 cultured pathogens only (*E. coli* and *Enterococcus* spp. respectively). In one case the Accelerate Pheno run failed on a sample harbouring *E. coli*, while BCID failed 3 times but when repeated was able to give correct results. Agreement of AST according to Accelerate Pheno and conventional testing was 97.5% (272/279 susceptibility tests performed). Overall, most disagreement of Accelerate Pheno with conventional testing was observed for amoxicillin-clavulanate susceptibility (3/19 cases), with Accelerate Pheno reporting as resistant 3 *E. coli* isolates testing as susceptible with Vitek 2. Time to results of different diagnostic tests are shown in Figure 2.

**Figure 2:**
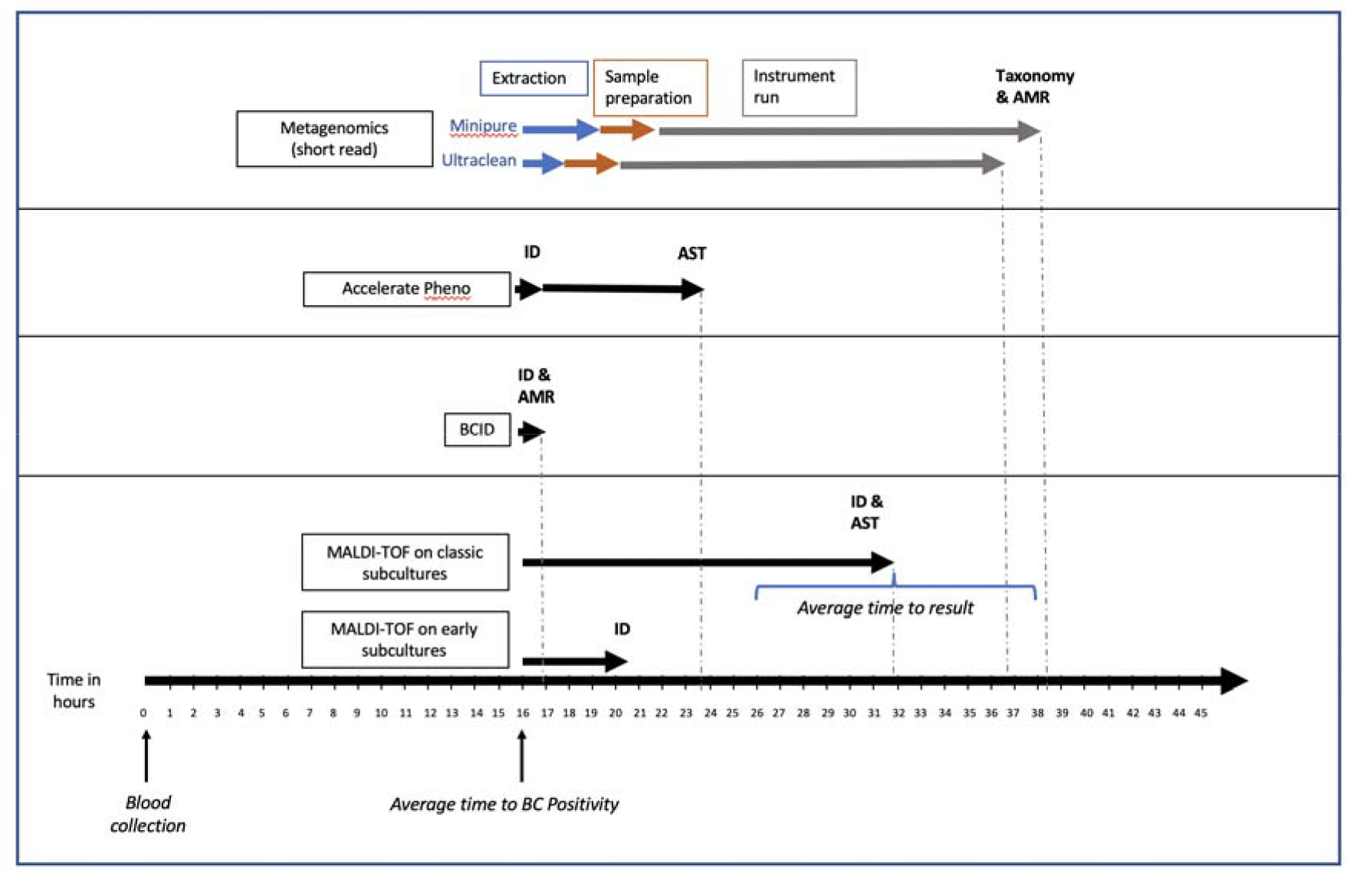
Time to results comparison of conventional culture, metagenomic WGS from positive blood culture broth (for short read) and commercial rapid diagnostic platforms

Average time to BC positivity for our samples was 16.1 h. Turnaround time of BCID for pathogen identification and antimicrobial resistance gene detection is 1 h from blood culture positivity while turnaround time of Accelerate Pheno is 1 h for pathogen identification and 7 h for AST. If implemented in a clinical laboratory, pathogen identification and WGS-AST based on short read direct metagenomic sequencing would be available approximately at the same time as results based on conventional testing. The time from fastq file upload to WGS-AST results via AREScloud is approximately 1 hour for metagenomic samples, including multiple samples run in parallel.

## DISCUSSION

A major barrier to direct sequencing from blood samples to detect pathogenic bacteria, is the limited sensitivity in patients with low loads of bacterial DNA in blood at the time of presentation ^29^. Adding a culture-amplification step by sequencing from positive blood culture broths leads to significant improvement in the amount of bacteria DNA available for sequencing. Currently, no protocols are available for host-depleted genomic DNA extraction direct from blood culture broths, which are also suitable for multiple downstream molecular applications. These often require removal of host genomic DNA and/or microbial elution at high yields. The MolYsis Complete and Basic kits effectively removed host DNA. The MolYsis Complete kit included host depletion and microbial extraction, however the DNA yield was too low for starting DNA input for long read sequencing, whereas Illumina’s Nextera Flex library prep permits an input as low as 1 ng/μL, with an adjustment to the amplification cycle step. The Basic kit uses host depletion only and is ideal to pellet microbial cells for alternative extractions methods. The subsequent extractions from both the Mini-pure and UltraClean methods had increased yields suitable for subsequent applications.

Previous published studies have utilized primarily spiked blood culture broths and extracted DNA without non-target human ‘host’ depletion ^3,30-33^. Host depletion is important for several reasons: there may be ethical restraints in the sequencing of human DNA; it mitigates against inefficiencies when over 80% of reads comprise off-target human DNA; and may optimise sensitivity for direct microbial DNA analysis. There have been a variety of commercial total DNA extraction kits reported for specific molecular assays. This study optimised a two-kit method with minimal out-of-kit modifications resulting in quality concentrated DNA for PCR and sequencing applications. The final DNA elution was host depleted, with removal of haem, SPS preservative (which acts as a PCR inhibitor) and other agents which neutralize antibiotics, whilst microbial DNA was concentrated, minimally sheared, and eluted in a non-EDTA buffer ^4,6,34^.

In an effort to ensure high DNA yields and options to increase low yield extractions, as well as remove short length DNA, kit protocol modifications or commercial kits were evaluated. The QIAamp Mini-pure kit was modified to concentrate DNA initially from a two-step 2x 50 μL to a 2x 25 μL elution with an optional SPRI bead cleanup eluting a smaller volume. This resulted in pure DNA at suitable concentrations for PCR and library preparation. The SPRI bead cleanup and Circulomics Short Read Eliminator protocols favoured size selection and successfully removed short fragments of DNA (∼4kbp) appropriate for long read sequencing to prevent fuel usage in flow cells, although short DNA removal may lead to the loss of resistance plasmids ^35^. DNA shearing by g-tubes technique produced DNA fragments suitable for sequencing long read ligation library preparations ^36,37^

Plasmid recovery from the extraction methods was assessed by downstream sequencing of a fully annotated reference genome (*E. coli* EC958) which contains two plasmids ^27^. The QIAamp Mini-pure enzyme-based extraction will not cleave plasmids and in turn, providing no ends for ligation library preparation ^35^. Although not undertaken, DNA shearing by g-tubes or megaruptor techniques produce DNA fragments suitable for sequencing library preparations ^36,37^. The UltraPure method using the mechanical and chemical lysis was effective at producing optimal DNA lengths with available ends. The resulting sequencing analysis correlated with published EC958 with an accurate number of plasmids identified.

The two extraction protocols optimised were effective for the removal of BC preservative and PCR inhibitors, SPS and other known inhibitors such as antibiotic neutralisers as verified by the PCR dilution experiments and purity data. The QIAGEN DNeasy Powerclean column was investigated and resulted in pure DNA however there was an observed consistent 50-80% loss of DNA yield, too low for long read sequencing. DNA preservative EDTA effects enzymatic activity during PCR and was replaced in the Mini kit to a non-EDTA buffer ^6,38^. The success of the pure, inhibitor free DNA was validated by PCR and observed no impact to DNA library preparation with the Illumina Nextera Flex transposase tagmentation process or Nanopore’s transposase or ligation methods.

The quality and purity of DNA was consistent with the optimised protocols, and additionally, with duration and storage temperature of BC. In the event a BC is unable to be extracted upon flagging positive, the duration and storage temperature experiments were investigated. The BCs were extracted on day 1 as the baseline, and every 24 hours up to day 5. Further, an aliquot of the BC was frozen at -80°C and extracted on day 5. In comparison to day 1, the quality and purity was maintained for each day and from the frozen extraction, short and long read sequencing analysis confirmed the identical AMR gene profile.

Correct species identification and detection of AMR genes from metagenomic data derived from clinical samples is a critical step in the application of direct sequencing for infectious disease diagnostics. However, correlation between the presence / absence of AMR genes and the resistance phenotype in order to guide appropriate antibiotic therapy, is not straight-forward ^39^. We employed a machine-learning algorithm, based on a sample bank with matched whole genome sequenced clinical isolates and AST results collected from several international centres ^13^. Our data show that phenotypic prediction from metagenomics data can be reliable for the most common Gram-negative pathogens encountered in patients presenting with bloodstream infection. However, training datasets will need to include a greater number of rarer pathogens or resistance phenotypes before reliability can be assured in these infrequent cases. The antibiotic agent for which WGS-AST was least reliable was ticarcillin-clavulanate, but this agent is not widely used in current practice (and is not commercially available in Australia, for instance). While the use of direct metagenomic sequencing and WGS-AST from positive blood culture broths holds promise, current methods using Illumina short-read sequencing remain time-consuming, and would offer few time advantages over conventional methods, and would be slower than other emerging rapid diagnostics, including those with predictive MICs (such as Accelerate Pheno). However, it is likely that other sequencing platforms, such as Oxford Nanopore, may be able to reduce the time to sequencing results and needs ongoing evaluation. The application of direct sequencing from blood cultures may also hold promise for the accelerated identification of slow growing, antibiotic affected or fastidious organisms, or where conventional phenotypic methods take days to complete.

Limitations to this study are acknowledged. We only used a limited number of samples and, while the range of organisms were prospectively collected and included common Gram-negative species, a broader range of pathogens, including diverse AMR phenotypes, would need to be assessed to understand the reliability of this approach. While we assessed the utility of DNA extraction methods for a variety of molecular applications, including long-read nanopore technology, we only used Illumina short read sequencing on all samples. While Illumina sequencing has high-fidelity, is a very reliable WGS method and is increasingly available in clinical laboratories, it can be slower than nanopore sequencing, which can return sequencing results in real-time. Further studies to reduce the time to results with nanopore sequencing from blood culture samples are warranted.

In summary, the UltraClean method proved optimal for host depleted, microbial enriched, inhibitor free DNA extraction and downstream molecular applications. Through one extraction, there is the ability to use DNA for PCR, short read and long read sequencing without plasmid loss. These methods support the development of molecular diagnostic assays and metagenomic sequencing direct from blood cultures, leveraging the advantages of pre-enrichment through culture amplification using commercial blood culture systems that are widespread in clinical practice. We have also demonstrated the utility of machine learning algorithms direct from clinical samples to accurately define effective antibiotic therapy. Further validation work and ultimately evaluation of the clinical benefit of such approaches are warranted.

## Acknowledgements and funding

This work (The “DETECT-ICU Project”) was funded by a grant from the Pathology Queensland Study, Education and Research Committee (SERC 5891_HarrisP) and the Royal Brisbane and Women’s Hospital Foundation. BCID and Accelerate Pheno kits were kindly provided by bioMérieux and Accelerate Diagnostics. PNAH is supported by an Early Career Fellowship from the National Health and Medical Research Council (GNT1157530).

