## Supplementary material for "An optimised method for bacterial nucleic acid extraction from positive blood culture broths for whole genome sequencing, resistance phenotype prediction and downstream molecular applications"

### Supplementary methods

Method 1 “Mini-Pure” Extraction: Microbial gDNA extraction commenced with the pellet being resuspended in 180µL of 20mg/mL lysozyme (Merck) and incubated at room temperature for 30 minutes. Genomic DNA was extracted manually and by QIAcube Platform using the QIAGEN QIAamp DNA Mini Kit following manufacturer’s instructions except for a two-step 2x 25µL elution with EB buffer (QIAGEN product 19086). Extractions were purified and or concentrated by SPRI Beads (Illumina). DNA shearing by g-TUBE (Covaris) for extractions used with ligation library preparation, was according to manufacturer’s instructions to attain 10 Kbp fragments by applying 6,000 RPM for 60 seconds.

Method 2 “UltraClean” Extraction: Microbial gDNA extraction commenced using the QIAGEN DNeasy UltraClean Microbial Kit with the pellet being resuspended in the 300 µL PowerBead solution, the suspended pellet transferred to a PowerBead tube and 50µL kit SL solution added. The tube was placed into a Labnet VorTemp 56 S205 and heat treated at 70°C for 10 minutes, thereafter the extraction followed manufacturer’s instructions.

### Supplementary Figures

Supplementary Figure S1. qPCR of the dilutions 1 in 10 to 1 in 10000, positive control blue, no-template control (NTC) orange. No inhibition for all dilutions.

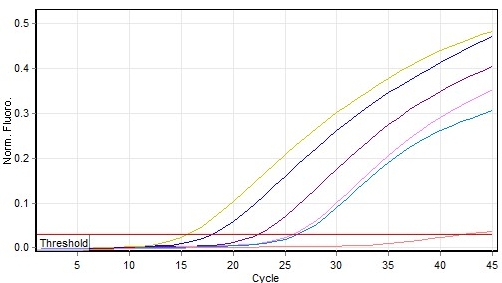

Supplementary Figure S2. A: ERV-3 qPCR for human DNA host depletion (below threshold), positive control blue, no human DNA present in extraction; B: 16S microbial DNA qPCR to confirm microbial DNA extracted from a fresh and frozen BC.

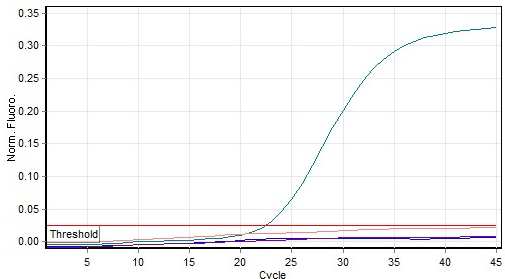

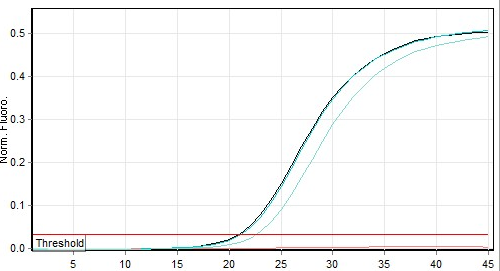

B

A

### Supplementary tables

Supplementary Table S1: Primer and probe sequences. Assay designed to detect Australian CMY variants (2, 4, 6, 16, 33, 42, 44, 48, 62, 68, 118 and 132).

| **Oligo Name** | **Oligonucleotide Sequence (5’-3’)** | **Designed by** |
| --- | --- | --- |
| Forward Primer | CTATCGCGAAGGGAAGC | This study |
| Reverse Primer | CATCTCCCAGCCTAATCC | Geyer *et. al.,* 2014,^1^ |
| Probe | 5HEX/CCGAAGCCTATGGCGTGAAA /3BHQ_1 | This study |

Supplementary Table S2: Sensitivity and Specificity testing of the *ampC* [*bla*_CMY-2-like_] real-time TaqMan PCR assay. Samples were extracted using boil prep method. Limit of detection (1 copy detected at Ct 36) of the assay was determined by digital droplet PCR from seven replicates of a 1 in 10000 positive CMY-2 control.

| **No. of Samples** | **Sample conc.** | **Positive for CMY** | **Negative for CMY** | **Sample Origin** |
| --- | --- | --- | --- | --- |
| 432 | Not specified | 32/32 | 400/400 | Harris *et al* 2018^2^ & this study |
| 20 | 1.0 McFarland | 10/10 | 10/10 | Harris *et al* 2018^2^ & this study |

Supplementary Table S3. Summary of extraction protocols and resulting DNA concentration, purity and length.

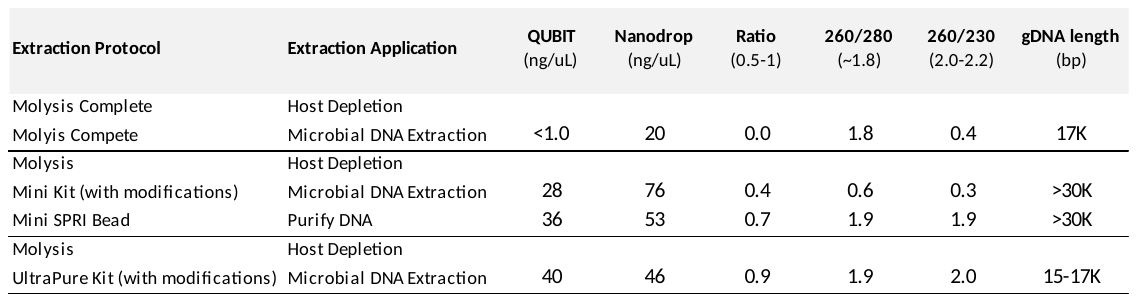

Supplementary Table S4: performance of WGS-AST for all species / antibiotic combinations

| **Taxon** | **Compound** | **CA** | **SN** | **SP** | **VME** | **ME** |
| --- | --- | --- | --- | --- | --- | --- |
| ***Escherichia coli*** | amikacin | 1.00 |  | 1.00 |  | 0.00 |
|  | amoxicillin and clavulanic acid | 0.86 | 1.00 | 0.80 | 0.00 | 0.20 |
|  | ampicillin | 1.00 | 1.00 | 1.00 | 0.00 | 0.00 |
|  | cefazolin | 0.64 | 1.00 | 0.62 | 0.00 | 0.39 |
|  | cefepime | 1.00 | 1.00 | 1.00 | 0.00 | 0.00 |
|  | cefoxitin | 0.93 |  | 0.93 |  | 0.07 |
|  | ceftazidime | 1.00 | 1.00 | 1.00 | 0.00 | 0.00 |
|  | ceftriaxone | 1.00 | 1.00 | 1.00 | 0.00 | 0.00 |
|  | ciprofloxacin | 1.00 | 1.00 | 1.00 | 0.00 | 0.00 |
|  | gentamicin | 1.00 | 1.00 | 1.00 | 0.00 | 0.00 |
|  | meropenem | 1.00 |  | 1.00 |  | 0.00 |
|  | norfloxacin | 1.00 | 1.00 | 1.00 | 0.00 | 0.00 |
|  | piperacillin and tazobactam | 1.00 |  | 1.00 |  | 0.00 |
|  | sulfamethoxazole and trimethoprim | 0.93 | 1.00 | 0.92 | 0.00 | 0.08 |
|  | tobramycin | 1.00 | 1.00 | 1.00 | 0.00 | 0.00 |
|  | trimethoprim | 1.00 | 1.00 | 1.00 | 0.00 | 0.00 |
| ***Klebsiella oxytoca*** | amikacin | 1.00 |  | 1.00 |  | 0.00 |
|  | amoxicillin and clavulanic acid | 0.00 | 0.00 |  | 1.00 |  |
|  | cefazolin | 1.00 | 1.00 |  | 0.00 |  |
|  | ceftazidime | 1.00 |  | 1.00 |  | 0.00 |
|  | ceftriaxone | 0.00 | 0.00 |  | 1.00 |  |
|  | ciprofloxacin | 1.00 |  | 1.00 |  | 0.00 |
|  | gentamicin | 1.00 |  | 1.00 |  | 0.00 |
|  | meropenem | 1.00 |  | 1.00 |  | 0.00 |
|  | piperacillin and tazobactam | 1.00 | 1.00 |  | 0.00 |  |
|  | sulfamethoxazole and trimethoprim | 0.00 | 0.00 |  | 1.00 |  |
|  | ticarcillin and clavulanic acid | 0.00 | 0.00 |  | 1.00 |  |
|  | tobramycin | 1.00 |  | 1.00 |  | 0.00 |
| ***Klebsiella pneumoniae*** | amikacin | 1.00 |  | 1.00 |  | 0.00 |
|  | amoxicillin and clavulanic acid | 1.00 |  | 1.00 |  | 0.00 |
|  | cefazolin | 1.00 |  | 1.00 |  | 0.00 |
|  | ceftazidime | 1.00 |  | 1.00 |  | 0.00 |
|  | ceftriaxone | 1.00 |  | 1.00 |  | 0.00 |
|  | ciprofloxacin | 0.50 |  | 0.50 |  | 0.50 |
|  | gentamicin | 1.00 |  | 1.00 |  | 0.00 |
|  | meropenem | 1.00 |  | 1.00 |  | 0.00 |
|  | piperacillin and tazobactam | 1.00 |  | 1.00 |  | 0.00 |
|  | sulfamethoxazole and trimethoprim | 1.00 |  | 1.00 |  | 0.00 |
|  | ticarcillin and clavulanic acid | 1.00 |  | 1.00 |  | 0.00 |
|  | tobramycin | 1.00 |  | 1.00 |  | 0.00 |
|  | trimethoprim | 1.00 |  | 1.00 |  | 0.00 |
| ***Proteus mirabilis*** | ampicillin | 1.00 | 1.00 | 1.00 | 0.00 | 0.00 |
|  | ceftriaxone | 1.00 |  | 1.00 |  | 0.00 |
|  | ciprofloxacin | 1.00 |  | 1.00 |  | 0.00 |
|  | gentamicin | 1.00 |  | 1.00 |  | 0.00 |
|  | sulfamethoxazole and trimethoprim | 1.00 | 1.00 | 1.00 | 0.00 | 0.00 |
|  | tobramycin | 1.00 |  | 1.00 |  | 0.00 |
| ***Pseudomonas aeruginosa*** | amikacin | 1.00 |  | 1.00 |  | 0.00 |
|  | cefepime | 1.00 |  | 1.00 |  | 0.00 |
|  | ceftazidime | 0.80 | 1.00 | 0.75 | 0.00 | 0.25 |
|  | ciprofloxacin | 1.00 |  | 1.00 |  | 0.00 |
|  | gentamicin | 1.00 |  | 1.00 |  | 0.00 |
|  | meropenem | 1.00 |  | 1.00 |  | 0.00 |
|  | tobramycin | 1.00 |  | 1.00 |  | 0.00 |
| ***Campylobacter jejuni*** | ciprofloxacin | 1.00 |  | 1.00 |  | 0.00 |

VME=very major error; ME = major error; TN=true negative; FP=false positive; FN=false negative; TP=true positive

Supplementary Table S5**.** Exploratory WGS-AST results using non-panel ResFinder 4 calls (based solely on presence of known AMR markers related to the compound in question, disregarding taxonomy)

| Species | Overall CA | Overall VME | Overall ME |
| --- | --- | --- | --- |
| All | 87.7 (402/458) | 50.1% (39/77) | 4.5% (17/381) |
| *C. jejuni* | 100% (2/2) | 0% (0/2) | 0% (0/0) |
| *E. coli* | 92.8% (219/236) | 30.3% (10/33) | 3.4% (7/203) |
| *K. oxytoca* | 68.8% (11/16) | 50% (4/8) | 12.5% (1/8) |
| *K. pneumoniae* | 96.9% (31/32) | 0% (0/2) | 3.3% (1/30) |
| *P. mirabilis* | 93.3% (42/45) | 75% (3/4) | 0% (0/41) |
| *P. aeruginosa* | 80% (40/50) | 90% (9/10) | 2.5% (1/40) |

WGS-AST = Whole genome sequencing antimicrobial susceptibility testing; CA = categorical agreement; VME = very major errors; ME = major errors; AMR = antimicrobial resistance genes
